# Patient-derived organoids assist personalized treatment for a patient with multiple metastatic neuroendocrine carcinoma of the cervix

**DOI:** 10.1101/2023.07.26.550503

**Authors:** Ruimei Liu, Xinyu Gao, Xiaoni Yue, Ying Yu, Zhaoting Xu, Feng Zhu, Leming Shi, Xinghua Cheng, Xinhua Lin, Yuanting Zheng

## Abstract

Neuroendocrine carcinoma of the cervix (NECC) is a rare subtype of cervical cancer with rapid metastasis. Lack of treatment regimens for metastatic NECC highlights the need for preclinical models and personalized treatment. Here, we established two patient-derived NECC organoid lines from lung metastases from the same patient, one of which was drug-free and the other exhibited tolerance to carboplatin plus etoposide (EP). The two organoids recapitulated the histopathology and genomic spectrum of the original tumors. Whole-genome sequencing revealed genomic structural variations (SVs), including HPV16 infection, NEUROD2, and ERBB2 amplification. The sensitivity phenotype of organoids to ERBB2 inhibitors coincides with the genomic features of ERBB2 amplification. In addition, the patient after relapse responded well to the paclitaxel plus carboplatin (TC) regimen as recommended by organoid drug screening, underscoring the utility of organoids as preclinical models. We first provide two NECC organoids of lung metastases with comprehensive molecular characteristics as valuable resources for this rare cancer and give a typical application for personalized treatment based on organoids.

## Introduction

Neuroendocrine neoplasias (NENs), characterized by neuroendocrine differentiation, are uncommon malignancies that arise from neuroendocrine cells, which are distributed widely throughout the body.^1,2^ NENs of the female genital tract are extremely rare with the cervix being the most common site.^3^ Neuroendocrine carcinoma of the cervix (NECC) is a poorly differentiated and aggressive cancer with a poor prognosis.^4^ Rapid metastasis is a typical behavior of NECC; the most common site of metastasis is the lung.^3^ Due to its rarity and metastasis nature, there are no standard treatment guidelines or preclinical models for NECC.^3,4^ Clinically, choosing appropriate treatment regimens and conducting prospective preclinical trials are challenging. Hence, there is an increasing need to better understand the biological characteristics of NECC to aid patients with this malignancy.

The common treatment for NECC is radical surgery combined with chemotherapy.^3^ Cisplatin/carboplatin plus etoposide (EP) is the most commonly used initial chemotherapy regimen.^5^ NECC exhibits diffuse growth and metastasis, resulting in a recurrence rate exceeding 50% within five years, and often develops resistance to EP chemotherapy regimens.^6^ For recurrent individuals, choosing effective treatment regimens is challenging and often fraught with uncertainty. To minimize the uncertainty and the side effects, personalized cancer treatment may be implemented. Personalized treatment based on better characterization of molecular and pharmacogenomic features of tumors aims to provide more effective treatments, which improves patient outcomes and reduce trials and error in the treatment process.^7^ For more accurate predictions of patients’ drug responses, personalized treatment is highly reliant on appropriate preclinical models.

Characterizing genomic profiles of preclinical models is key for personalized treatment. For mutational variants, alterations in TP53 and RB1 proved common in neuroendocrine carcinomas.^8,9^ In contrast to the other neuroendocrine carcinomas, TP53, KRAS, PIK3CA, and KMT2C are the most frequently found mutations in NECC.^10^ Moreover, structural variations (SVs) play a key role as connections or rearrangements between remote genomic loci and underlie all copy number alterations (SCNAs), which may have a vital effect on NENs.^8,9,11^ Additionally, NECC is associated with the human papillomavirus (HPV), especially HPV16 and HPV18, which are involved in the progression of neuroendocrine carcinoma.^12,13^ These genetic backgrounds may underlie molecular mechanisms that contribute to finding therapeutic targets. However, the gene-drug association treatment strategy has been hindered by the paucity of biological understanding of tumor response to drugs.^14^ Therefore, establishing preclinical models and comprehensively characterizing them is essential for ensuring the safety and efficacy of therapies and for advancing our understanding of NECC and its treatments.

Recently, patient-derived organoids (PDOs) have become reliable preclinical models.^7,15^ PDOs recapitulate many structural and functional characteristics of their in vivo corresponding organs.^16^ It has, to some extent, circumvented the problems of genetic drift in cell line passages and the difficulty of constructing xenograft mouse models for a long time.^16,17^ Organoids of gastroenteropancreatic neuroendocrine neoplasm and common cervical cancer patient-derived organoids have been established, which play a vital role in the comprehension of corresponding cancers and personalized medicine.^11,18^ However, NECC is not included in either of these biobanks. Due to some unique characteristics of NECC, like HPV infection, the developed culture system may not be suitable for NECC. It is still a challenge to develop organoid models for this rare cancer.

To take one small step further, a NECC patient with two lung metastases was enrolled in this study. Using a modified protocol, we successfully established two patient-derived organoid lines from two lung metastases in the same NECC patient and conducted a comprehensive genomic characterization of these two organoids. Moreover, we verified the validity of the organoids as pharmacological screening models. Notably, we have used these two organoid lines to guide the chemotherapy process, and this NECC patient had potentially benefited from it. Our study not only provided two valuable organoid lines with comprehensive genome and transcriptome features as preclinical research models for NECC, but also served as an organoid-based precision medicine case for this rare cancer.

## Results

### Overview of the study

A NECC patient with two lung metastases was enrolled in this prospective study. We have followed this patient over a period of three years and successfully established two NECC organoid lines derived from the two successive lung metastases (Figure 1A). To confirm the origin and comprehensively characterize the two organoids, we performed whole genome sequencing (WGS) and RNA sequencing (RNA-seq) on the organoids and the corresponding formalin-fixed paraffin-embedded (FFPE) tumor tissues (Figure 1B). WGS and RNA-seq were technically repeated twice to reduce the artifacts (Figure 1B).^19,20^ The intersection of the two WGS technical replicates was used to identify SNVs, Indels, and SVs across the whole genome and exon region, respectively (Figures S1A, S1B). Similarly, gene expression profiles were used by taking the averaged FPKM values of the two technical replicates from RNA-seq (Figures S1C, S1D). Based on the molecular and pathophysiological characteristics, personalized chemotherapy regimen was elaborated according to the drug response of the organoids (Figure 1C).

**Figure 1.**
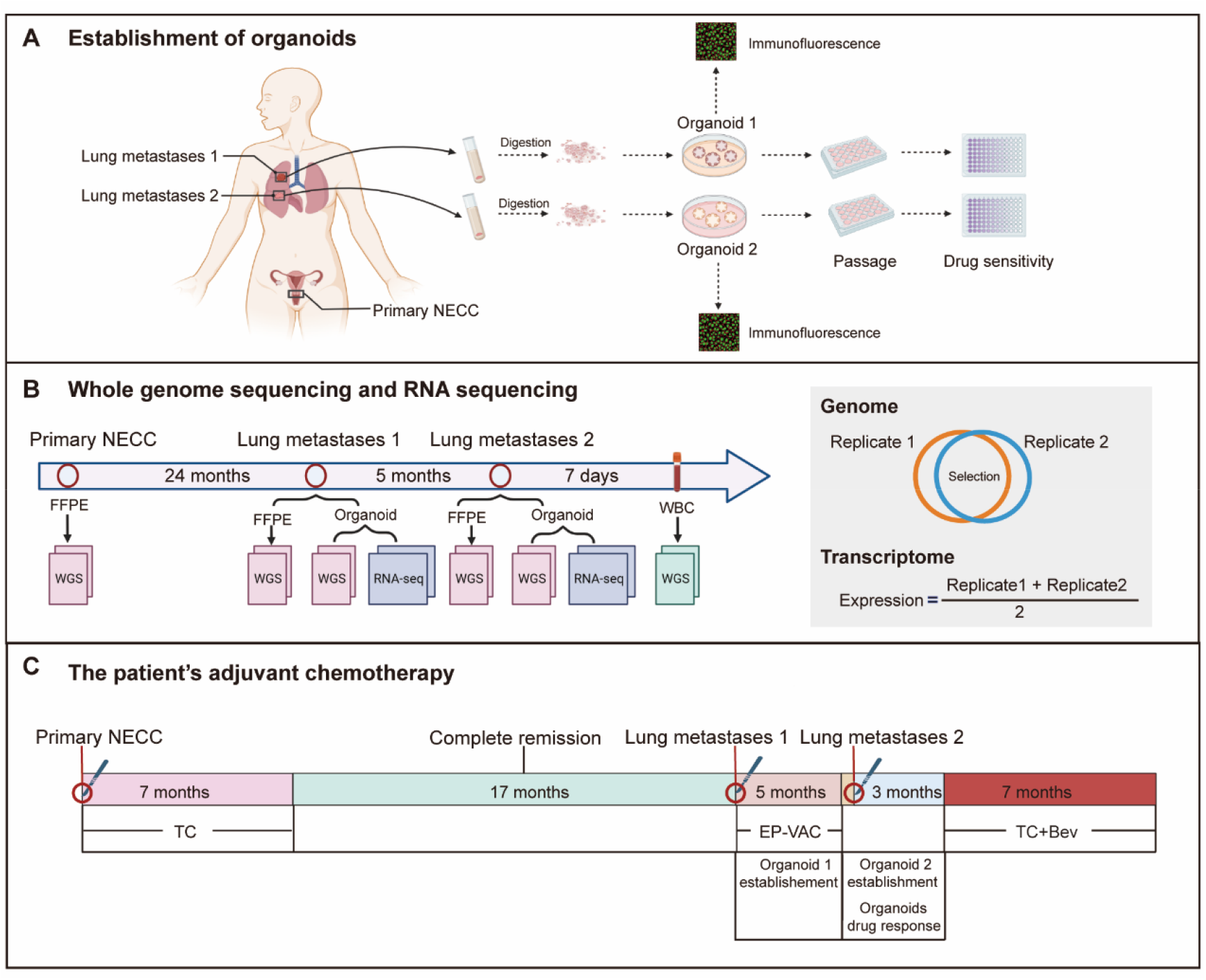
Overview of the study. (A) The whole process of establishing organoids. Two organoid lines were established from two lung metastases of the same NECC patient. Immunofluorescence was used to determine the origin of the two organoid lines. Drug sensitivity was tested after stable organoid passage. (B) Whole genome sequencing was performed on tumor tissues and corresponding organoids; RNA sequencing was performed on organoids; WBC means white blood cell. To get the genomics and transcriptomics profiles of each sample, two replicates were sequenced. (C) The treatment regimen and time course of the NECC patient from primary to two metastases.

### Establishment of patient-derived NECC organoid lines from two successive lung metastases

We aimed to establish two clinically relevant metastatic NECC organoid lines that were amenable to drug screening. We collected two successive metastases to the lung from the same NECC patient. Given pulmonary colonization and characteristics of NECC, we optimized the organoid culture system of lung cancer and cervical cancer to develop our final culture system.^18,21-23^ Initially, irregular and cauliflower-shape organoids were formed using the conventional lung organoid conditional culture medium system. However, organoid growth and maintenance were still limited under these basic conditions. Therefore, we added epithelial growth factor (EGF) combined with basic key factors such as R-Spondin1, Noggin, B27, and fibroblast growth factor (FGF) to ameliorate the culture conditions, which obviously improved the proliferation rate and the state of organoids (Figure 2A, Figure S2A). By adopting the optimized protocol, the second metastasis organoid was also successfully established, further demonstrating the effectiveness of the culture system. The morphology of the two organoid lines was stable and similar throughout the cell passage process (Figure 2B, Figure S2B). Both of them were stably cultured for long-term expansion over six months. Until now, the two organoids have been cultured for 1.5 years and still exhibit good vitality (Figure 2SB). During the cultivation, we observed that the second organoid exhibited a slightly slower rate of proliferation compared to the first one (Figures S2B and S2C). To explain this phenomenon, we assessed the differences in gene expressional patterns of the two organoids using the RNA-seq data. All detected genes were highly similar, with a Jaccard similarity coefficient of 87.6% between the two organoids (Figure S2D). Compared to the first metastasis organoid, the up-regulated genes of the second were enriched in the pathway of response to virus, indicating it may be associated with HPV infection (Figure S2E). Compared to the second metastasis organoid, five cancer-related genes associated with cell proliferation and differentiation were considerably upregulated in the first metastatic organoid, which may explain the faster proliferation rate of the first lung metastasis organoid (Figures S2F and S2G). Taken together, the results demonstrated the efficiency of our NECC organoid culture protocol and explained the distinction in growth states of the two organoids at the transcriptional level.

**Figure 2.**
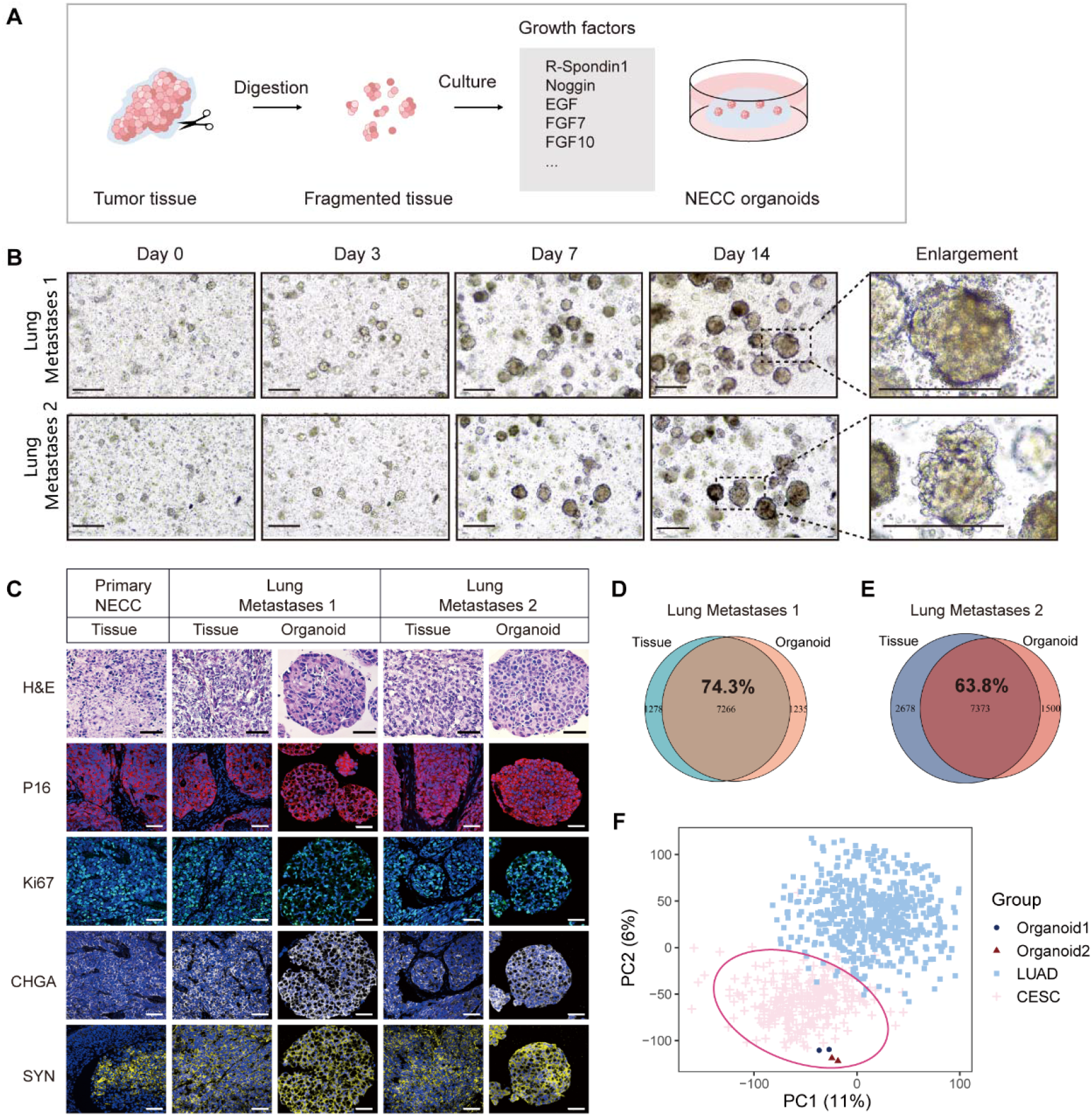
Establishment and characteristics of the two patient-derived metastatic NECC organoids. (A) Schematic overview of establishing NECC organoids. (B) Representative images during organoid culture on days 0, 3, 7, and 14; scale bar indicates 300 μm. (C) Hematoxylin and eosin (H&E) and immunofluorescent staining of HPV (P16), cell proliferation (Ki67) and NECC markers (SYN and CHGA) for tumor tissues and established organoids. Scale bar indicates 50 μm. (D, E) Consistency of SNVs and Indels between tumor tissue and the established organoid in whole genome. (F) In the principal component analysis (PCA) plot of gene expression, the two organoids were grouped into cervical squamous cell carcinoma, endocervical adenocarcinoma (CESC), but not lung adenocarcinoma (LUAD).

### Organoids recapitulate the histological and genetic characteristics of the original cancer tissues

It is crucial that the established organoids sufficiently represent the diverse histological and genomic features of the original tumor tissue. To trace back the origin of the two organoid lines, we analyzed Hematoxylin and eosin (H&E) staining and the expression of Chromogranin A (CHGA), Synaptophysin (SYN), and P16 by immunofluorescence staining (IF).^24^ P16 associated with high-risk human papilloma virus (HPV) infection and Ki67 (>60%) were positive, suggesting high cell proliferation and poor prognosis (Figure 2C). In addition, the NECC-related markers, including CHGA and SYN, were positive (Figure 2C). These results displayed that the NECC-related markers were positive and the same staining patterns were exhibited between original tumor and the corresponding organoid (Figure 2C). To check whether the organoids preserved the genetic alterations of the original cancer tissues, we performed WGS analysis on both of them. Genome-wide somatic mutations were highly consistent between organoids and the original tissue, with a Jaccard similarity coefficient of 74% and 64%, respectively (Figures 2D, 2E). Given that the metastatic NECC tissue was taken from the lung, we further explored whether the organoid source was lung or NECC. We integrated gene expression data of lung adenocarcinoma (LUAD), cervical squamous cell carcinoma and endocervical adenocarcinoma (CESC) and the two organoids. As shown in the PCA plot, the NECC organoids were clustered with CESC, which provided further evidence that the metastasis tumor was from the primary NECC (Figure 2F). All together, these results indicated that the patient-derived NECC organoids faithfully recapitulated the NECC tissues and maintained their genetic status and phenotypes during long-term expansion in vitro.

### Mutational signatures showed similarity between tumor tissues and organoids, and an increasing trend of mutation burden from primary NECC to lung metastases

To profile the mutation status and evolution during the whole course of NECC, we detected somatic mutations (SNVs and Indels) in whole exon regions and selected sense mutation type including nonsynonymous, frame shift insertion, frame shift deletion and stop gain (Figure 3A, Table S1). NECC organoids retain most sense mutations from the original tissue, with Jaccard similarity coefficients of 87% and 82%, respectively (Figure 3B). The tumor mutation burden value was about two, which was similar to the common neuroendocrine carcinoma.^8^ Furthermore, metastases exhibited a higher mutation burden than that of the primary tumor tissue (Figure 3C). Next, we selected cancer-related genes reported by TCGA and NECC-related genes in COSMIC.^25^ Seven cancer-related genes were detected in both primary tumor and metastases, among which KMT2C and ERBB2 were also detected in other NECC studies (Figure 3D).^8,9^ Novel mutations in three cancer-related gene were detected only in the metastases, including FAT1, FGFR2 and CIITA, which may be related to metastasis and resistance to therapy (Figure 3D).^26,27^ The variant allele fraction of these cancer-related genes was mostly higher in organoids than that in the matched tumor tissues. (Figure 3E). To further characterize the genome-wide mutational profile, we performed mutational signature analyses and found the APOBEC signature which mediated antiviral response was present in both NECC tumor tissues and matched organoids (Figure 3F). APOBEC activity may continually contribute to mutagenesis during tumor progression.^28,29^ Simultaneously, evolutionary behaviors showed the parallel evolution from primary to recurrent metastasis (Figure S3). Taken together, the organoids maintain key mutation features of NECC tumor tissues and limited evolution was shown during the progression.

**Figure 3.**
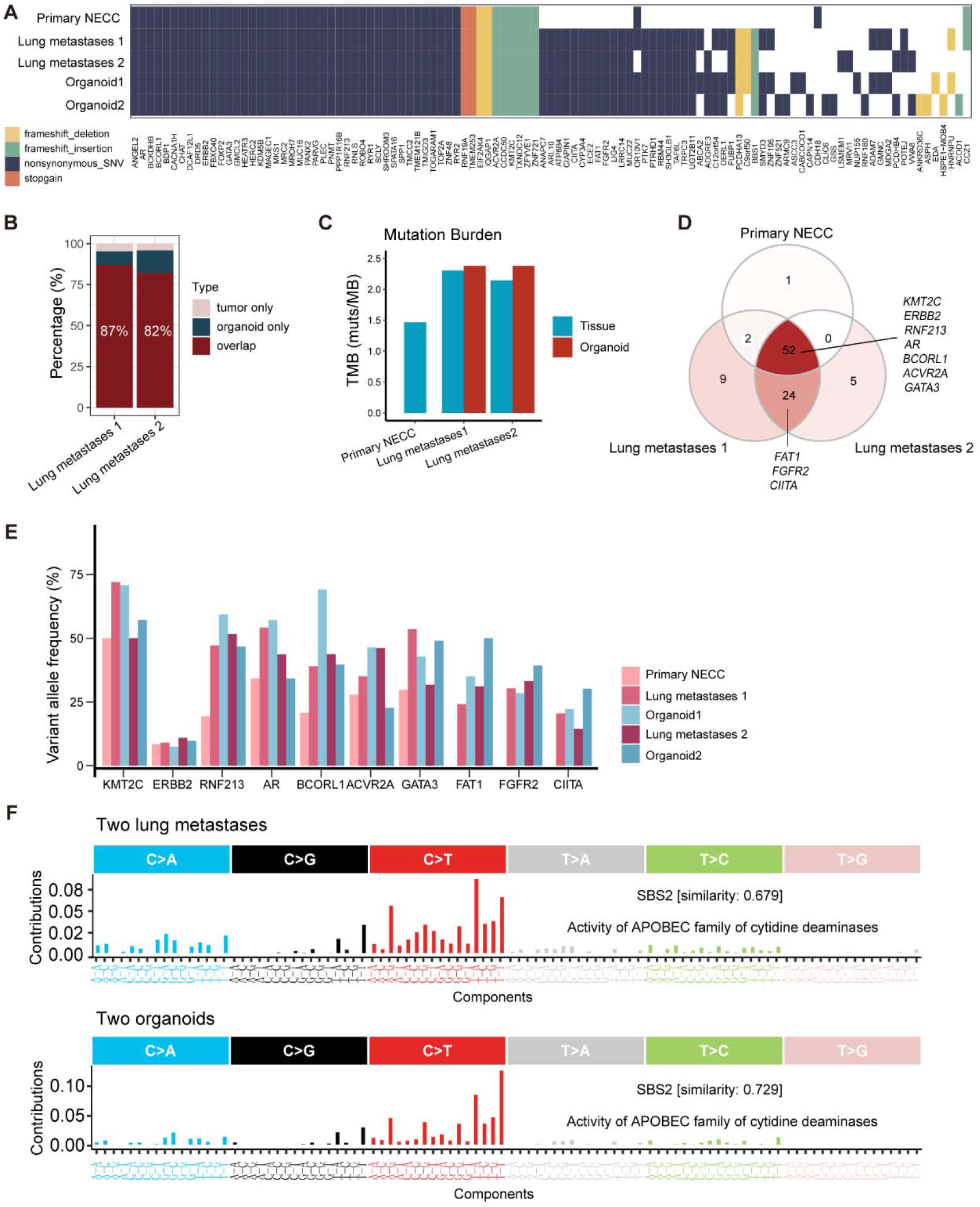
Genetic mutations profiles showed similarity between tumor tissues and organoids. (A) The genetic landscape of tumor tissues and organoids in gene coding region (B) Concordance of somatic mutation between tumor tissue and its derived organoid in the whole exon region (C) Tumor mutation burden of tumor tissue and organoid (D) Venn diagram shows the regions of shared and unique mutations among the primary NECC and recurrent lung metastasis tumor tissues. (E) Variant allele frequency of the cancer-related genes of tumor tissue and organoids. (F) Mutational signature analysis of two metastases (up) and corresponding two organoids (down) displayed the same pattern.

### Structural variations and HPV16 infection exhibit similar pattern between tumor tissues and organoids

In order to characterize SVs and HPV infection from primary to metastatic stages, we performed WGS analysis on NECC primary tumor tissues and corresponding organoids. The organoids displayed the same SV pattern as the parental cancer tissues, indicating that they retained the SV signatures of the matched cancer tissue (Figure 4A). SVs were more prevalent in lung metastases than the NECC primary (Figures 4A, 4B). In addition, the deletion events account for the highest percentage of all SV types, the percentage of which was higher in primary NECC than recurrence metastases (Figure 4B). The high percentage of SV deletion may cause the NECC progression in this patient.^30^ We did the profiles of HPV insertion sites, indicating that all samples were infected with HPV16 (Figure 4C, Figure S4, Table S4). The insertion sites of HPV16 were mainly concentrated on chromosome 17, 11, 1 and 3 (Figure S4). Next, we detected the shared HPV16 insertion sites in both NECC primary and metastases (Figure 4D). The shared HPV16 insertion sites were located in the region of ERBB2, NEUROD2, BARX2 and HNF1B (Figure 4E). ERBB2, BARX2 and HNF1B were related to the progression and invasiveness of cancer.^31-33^ In particular, HPV16 was inserted in the 3’UTR region of the NEUROD2 gene in all samples, which may lead to elevated expression of NEUROD2 related to the neuroendocrine phenotype (Figure 4E). Taken together, significantly recurrent SVs and HPV16 insertions may be contributing factors to the occurrence and progression of NECC in this patient.

**Figure 4.**
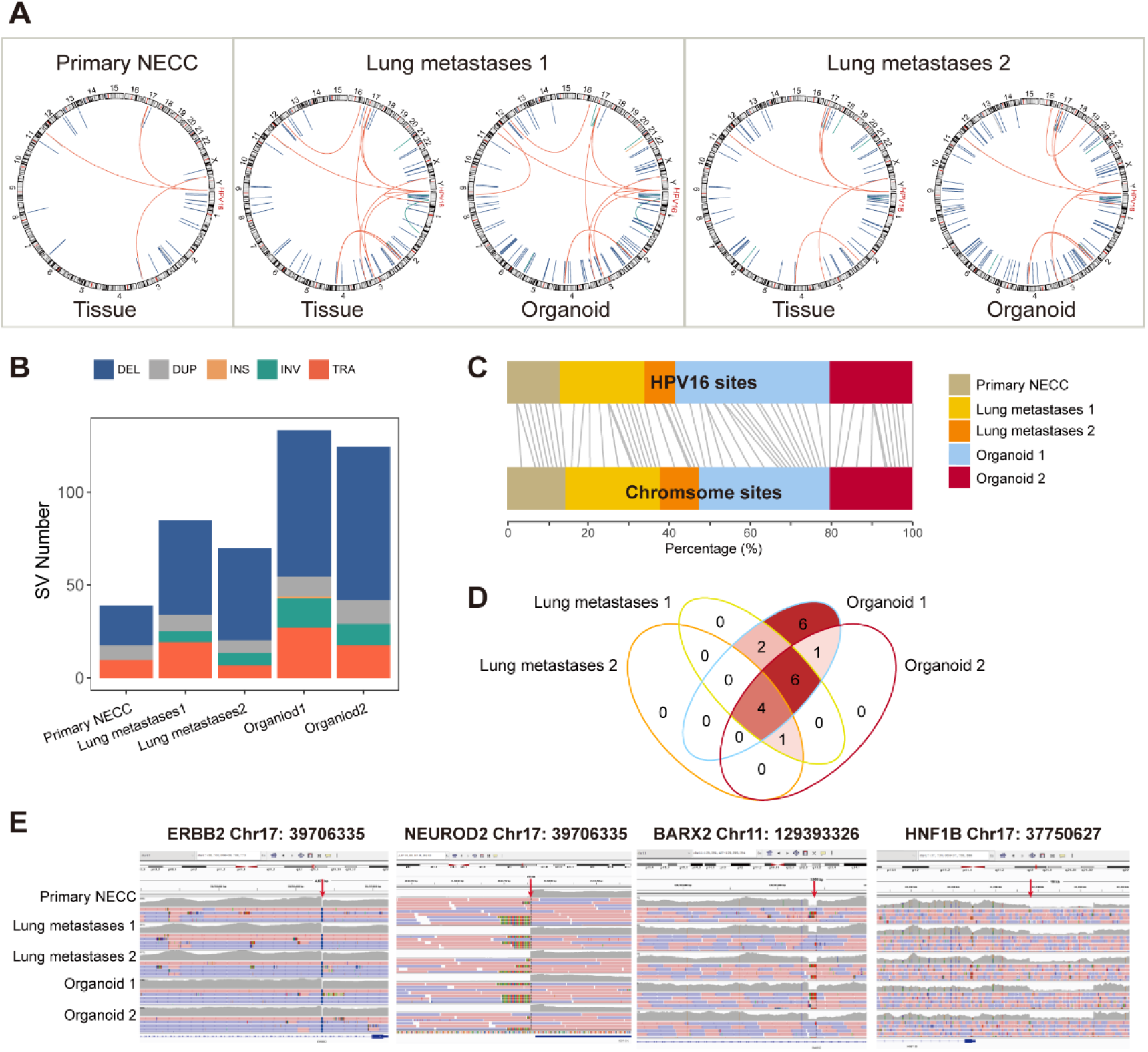
Structural variations and HPV16 infection exhibit recurrence from primary NECC to lung metastases and corresponding organoids. (A) SV patterns from primary NECC to lung metastases and corresponding organoids (B) Number of SV types (DEL: deletion; DUP: duplication; INS: insertion; INV: inversion; TRA: translocation) in tumor tissue and corresponding organoid. (C) The number of HPV16 insertion sites in the five samples, including tumor tissue and organoids. (D) Venn diagram showing the numbers of shared and unique HPV16 insertion sites. (E) IGV (Integrative Genomics Viewer) showing details about shared HPV16 insertion sites related to cancer genes.

### High Copy number and overexpression of ERBB2 and NEUROD2 show recurrence from primary NECC to lung metastases and corresponding organoids

Copy number alterations analysis revealed a greater variety of chromosomal changes in metastatic tissues and a high concordance between primary tissue and the matched organoids (Figure 5A). Chromosome 17 showed a high instability in both primary and metastatic samples (Figure 5A). According to the gain and loss list (Table S3), we selected the cancer related genes and detected recurrent amplification of ERBB2, ETV5 and recurrent deletion of TP53, GAS7, USP6 and RABEP1. ERBB2 exhibited absolute copy-numbers (CNs) of 90 in primary tissue and more than 120 in metastases. (Figure 5B). Furthermore, we checked the CNVs of the NEUROD family genes related to the neuroendocrine phenotype. NEUROD2 exhibited a high absolute CNs of six in primary tissue and more than 17 in metastases (Figure 5B).

**Figure 5.**
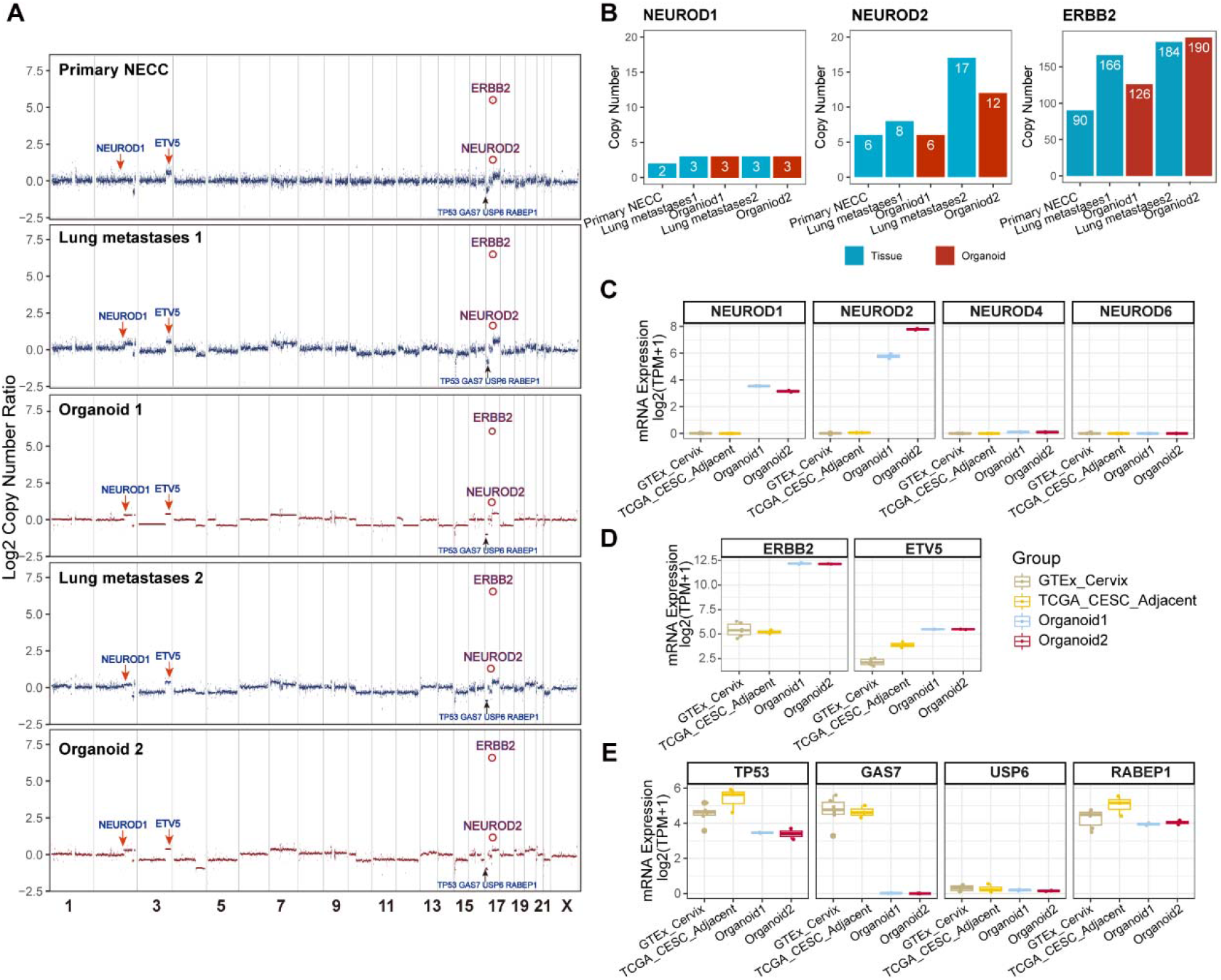
Copy number profiles coincide with gene expression levels from primary NECC to lung metastases and corresponding organoids. (A) The copy number alerts status of tumor tissue and the corresponding organoids. (B) The absolute copy number of NERUOD1, NERUOD2 and ERBB2. (C) RNA expression level of genes related to NEUROD family genes. RNA expression [log2(TPM+1)] of organoids, the normal cervix tissue expression data downloaded from GTEx and CESC adjacent tissue expression data downloaded from TCGA. (D) RNA expression level of genes related to the gain of genes in tissue and the organoids. (E) RNA expression level of genes related to the loss of genes in tissue and the organoids.

We next validated our findings by using the RNA-seq data to detect expressional level of these genes. Genes with copy number gain have higher expression and genes with copy number deletion have lower expression than the normal samples of GTEx cervical samples and CESC adjacent samples from the TCGA database (Figures 5C, 5D and 5E; Table S5). Among the NEUROD family, NEUROD2 and NEUROD1 with higher CNs showed higher expression than NEUROD4 and NEUROD6 (Figure 5C). Research has proven that NEUROD2, a member of the NEUROD family of neurogenic basic helix-loop-helix (bHLH), regulates neuronal migration.^34,35^ So the high copy numbers and high expression of NEUROD2 may be associated with the metastasis-prone phenotype of NECC. Furthermore, TP53, a tumor suppressor gene, was deleted and exhibited low expression, which may also contribute to the metastasis-prone phenotype (Figure 5E). ERBB2, which encodes a member of the EGF receptor family of receptor tyrosine kinases, is particularly highly expressed (Figure 5D). This suggests that ERBB2 may be an effective therapeutic target for this patient.^36^ Taken together, chromosomal changes indicated that the genome was more instable after recurrent metastases, with a subtle difference between the first and the second metastases.

### The patient’s response to TC regimen after relapse is consistent with the drug response of patient-derived organoid lines

To assess the utility of the two organoid lines as preclinical models for drug response evaluation, we performed dose titration assays using 11 compounds and three multi-drug combination chemotherapy regimens. The two organoid lines were in passage eight and displaying phenotypic stability. Drugs were selected based on this NECC patient’s clinical treatment, NCCN (National Comprehensive Cancer Network) recommendations and targeted therapeutics, including ERBB2 inhibitors, anti-mitotic agents, topoisomerase inhibitors and DNA synthesis inhibitors, and DNA alkylators (Figure 6A). To assess the drug sensitivity of organoids as preclinical models for drug testing, we defined the drug sensitivity interval in vivo according to the pharmacokinetic Cmax data of phase I clinical trials of these compounds (Figure 6B).^37-44^

**Figure 6.**
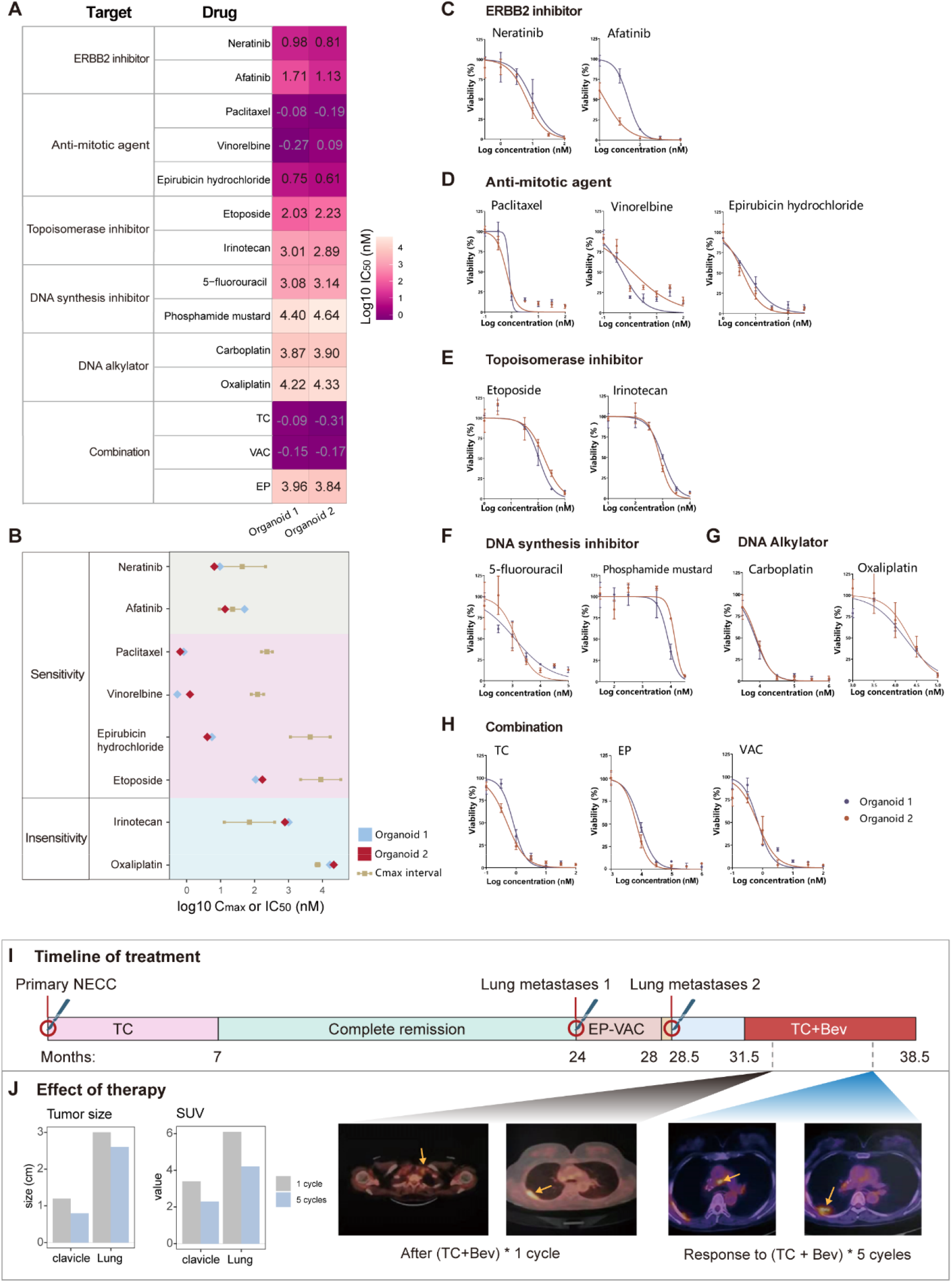
Drug response and clinical utility of the two organoid lines. (A) IC50 values were calculated from drug response analyses of two patient-derived organoids by applying nonlinear regression (curve fit) after different drug treatments. (B) Comparison of IC50 values and the maximum drug concentration (Cmax) of the same drugs. IC50 ≤ Cmax was defined as sensitivity, while IC50 > Cmax was defined as insensitivity. Range bars represent Cmax. (C-H) Dose response curves for the two NECC organoids treated with different drug or drug combination. TC: paclitaxel plus carboplatin; EP: carboplatin plus etoposide; VAC: vinorelbine, phosphoramide nitrogen mustard plus epirubicin hydrochloride (I) Timeline of treatment from primary to two recurrent metastases. EP-VAC means EP and VAC were used interchangeably. TC + Bev means TC and Betovazumab in combination. (J) Tumor size and standard uptake value (SUV) were reduced from 1 cycle to 5 cycles of TC+Bev. On the right are positron emission tomography and computed tomography (PET-CT) images of the lung and clavicle after one cycle of the TC plus Bev regimen and PET-CT images of the lung and clavicle after another four cycles of the TC plus Bev regimen.

The two organoid lines displayed similar drug responses to the selected compounds (Figures 6A and 6B). For the ERBB2 amplification target, the two organoid lines displayed significant responses to neratinib and afatinib (ERBB2 inhibitors). The second organoid showed greater sensitivity to ERBB2 inhibitors, especially afatinib (Figure 6C), as it had higher ERBB2 amplification than the first one (Figures 6B, 6C). Both organoids exhibited drug sensitivity to anti-mitotic agents and topoisomerase inhibitors, including paclitaxel, vinorelbine, epirubicin hydrochloride, etoposide and irinotecan (Figure 6B). However, these chemotherapy drugs did not kill the organoid thoroughly even at high concentrations (Figures 6D, 6E). Notably, paclitaxel showed a strong killing effect even at very low concentrations (Figure 6D), suggesting its potential as an effective choice for combination chemotherapy. The two organoids showed insensitivity to DNA synthesis inhibitors, including 5-fluorouracil and phosphamide mustard (Figure 6F) and displayed drug sensitivity to carboplatin but not to oxaliplatin (Figure 6G). Significantly, carboplatin had a completely organoid-killing effect and played a key role in combination chemotherapy. Then, we examined the effects of the selected combination chemotherapy regimen including paclitaxel plus carboplatin (TC), carboplatin plus etoposide (EP) and vinorelbine, phosphoramide nitrogen mustard plus epirubicin hydrochloride (VAC) (Figure 6G). Consistent with the effect of the single compound, TC worked best, followed by EP and VAC regimens.

Based on the drug response of the two patient-derived organoid lines, this NECC patient was considered to potentially benefit from the modified chemotherapy regimen. After receiving a first-line TC regimen, this NECC patient had a clinically effective reaction and kept complete remission for 17 months (Figure 6I). EP and VAC were used alternately for four cycles after the first recurrence surgery (Figure 6I). But her tumor did not respond to this regimen and recurred during the course of treatment (Figure 6J). As a result, the patient underwent a third resection. In contrast to EP and VAC, the second organoid showed a more significant drug response to the TC regimen. Based on this, the chemotherapy was changed from EP and VAC to TC plus bevacizumab (Bev), which had a positive effect (Figures 6I, 6J). After another four cycles of TC plus Bev treatment, the patient’s lung tumor decreased in size from 3.0 cm to 2.6 cm, and the lymph node metastasis of the clavicle decreased from 1.2 cm to 0.8 cm (Figure 6J). This indicates a strong consistency between the drug response of the patient-derived organoids and the drug response of the patient’s cancer in vivo. Additionally, both of the two organoids showed a positive response to ERBB2 inhibitors. Based on the drug response, TC combined with an ERBB2 inhibitor might be the best selection. Unfortunately, due to the patient’s medical condition, ERBB2 inhibitors were not used. These findings suggest that the two organoids could be appropriate preclinical models for testing the drug response of this NECC patient and providing some evidence for prospective personalized treatment.

## DISCUSSION

Although NECC is a rare tumor, its incidence has been increasing, which may be attributed to the progress of diagnostic technology accompanied by the development of precision medicine. To our knowledge, no preclinical organoid models are available, which has been an obstacle for the NECC study. Here, we established and characterized two NECC lung metastasis organoid lines. To provide comprehensive and accurate molecular information, we analyzed two replicates for WGS and RNA-seq to minimize the false positive sites. Notably, the two organoids are not isolated individuals but have internal connections and unique characteristics. The first organoid was a treatment-naive organoid model, and the second was a drug-resistant model of the EP-VAC chemotherapy regimen, which is an effective model for studying EP-resistant patients. We provided two valuable preclinical model for this field.

Our study demonstrates that patient-derived NECC organoids showed practical utility for personalized medicine. Characterizing molecular profiles in combination with therapeutic compounds can reveal druggable targets or suitable chemotherapy regimens. In this study, we examined common chemotherapy regimens and targeted drugs on the two organoid lines. For the druggable target, we found that ERBB2 had high copy numbers and high expression in both organoids and the corresponding tumor tissue, and organoids showed higher sensitivity to ERBB2 inhibitors. We predicted that TC plus ERBB2 inhibitors might have effective effect on this case. Also, molecular profiles showed that the two metastases retained most of the features of the primary cancer, which suggests that the TC regimen might still be effective for the second metastatic cancer. Meanwhile, the two organoids indeed still manifested drug response to the TC regimen. Based on the drug response, the patient’s chemotherapy regimen was adjusted to TC plus Bev regimen. The overall survival for this NECC patient was significantly longer than the median overall survival of patients with this deadly disease, which suggests a potential benefit for this case. For more precise and secure chemotherapy, the drug effect uncertainties can be partly eliminated based on the drug response of the NECC organoids.

Interestingly, we found that NEUROD2 had high copy numbers and high expression from primary NECC to metastases. NEUROD2 amplification may be a key factor in tumor differentiation toward a neuroendocrine phenotype.^34,35^ Gain of this phenotype may be a factor driving metastases of NECC. Suppressing the expression of this gene may provide some clues for the treatment of the NECC metastasis. Further investigation is required to delineate molecular mechanisms of this phenomenon.

Some questions still remain: For example, the sample size of this study was limited, and the scarcity of patients with NECC is a barrier for establishing organoid biobank with a large number of NECC patients. Next, more samples will be collected to test the culture environment for establishing NECC organoid and to provide a more comprehensive molecular profile for ensuring the clinical utility. Multicenter consortia may be a solution for preclinical model construction of rare cancers like NECC.

In conclusion, this study highlights the clinical usefulness of NECC organoids as an effective preclinical model. From primary to metastatic NECC, the enhancement of tumor aggressiveness and metastatic ability is accompanied by a series of changes in the genome, including the increase of the number of SVs, HPV integration, and gene copy number of some cancer related genes, such as NEUROD1, NEUROD2, and ERBB2. The two organoids recapitulate the histopathological and molecular spectrum of the corresponding tumor tissues. This study served as an effective practice to bridge the gap from bench to bedside.

## ACKNOWLEDGMENTS

We thank the patient who consented to donate her tumor tissues for this study, as well as the surgical team.

## AUTHOR CONTRIBUTIONS

Y.Z., R.L., X.L. designed the study. XG., R.L. and XZ generated and cultured organoid lines. RL performed the bioinformatic analyses and data interpretation. X.G. performed H&E staining, immunostaining, and drug response assays. X.C. provided tumor tissue samples. R.L, X.Y. collated clinical information and helped with data interpretation. Y.Y., L.S. and F.Z. helped checked out manuscript. RL prepared the manuscript with help from all authors. Y.Z, X.L., X.C., conceived the project and helped with data interpretation.

## STAR Methods

### Key resources table

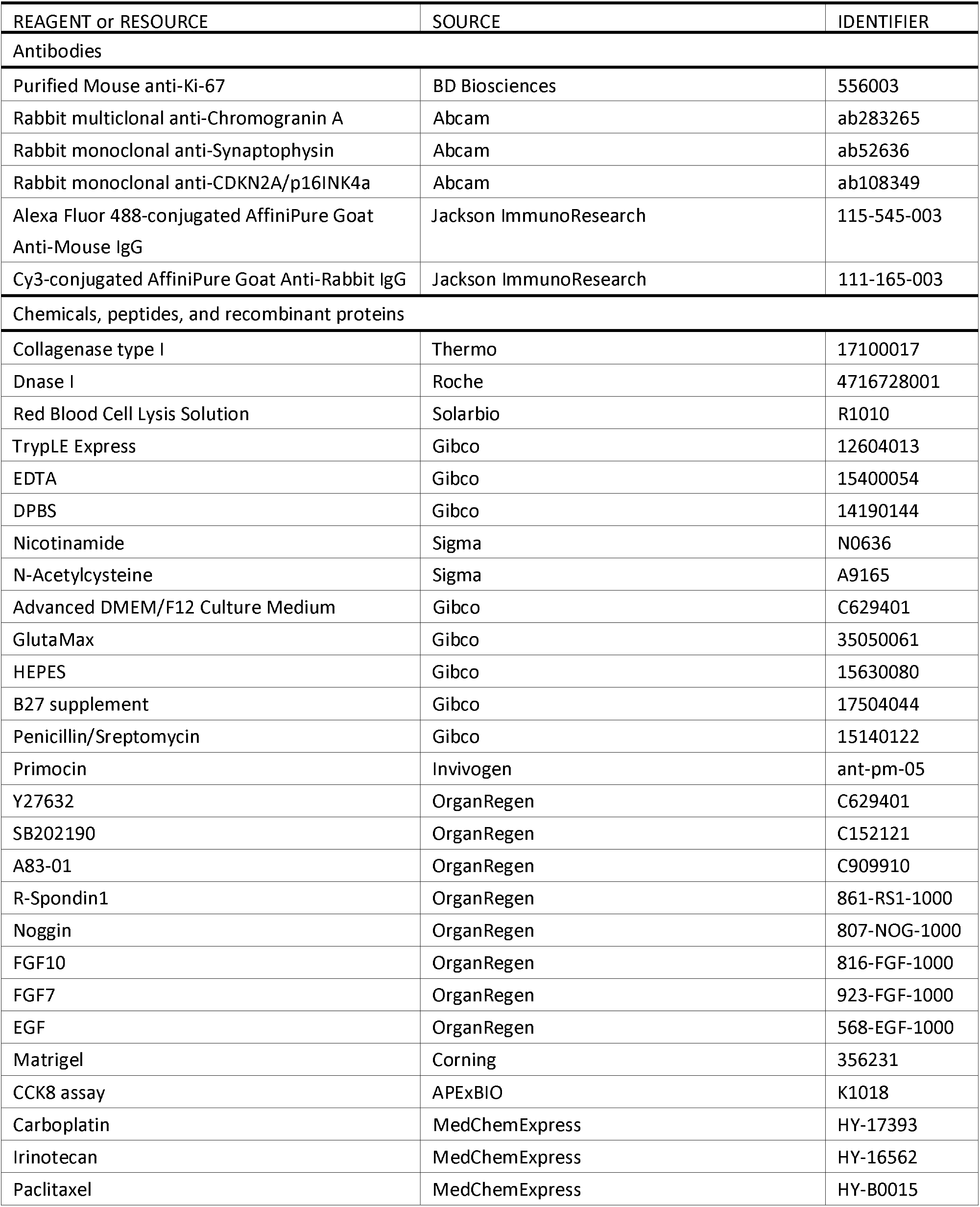
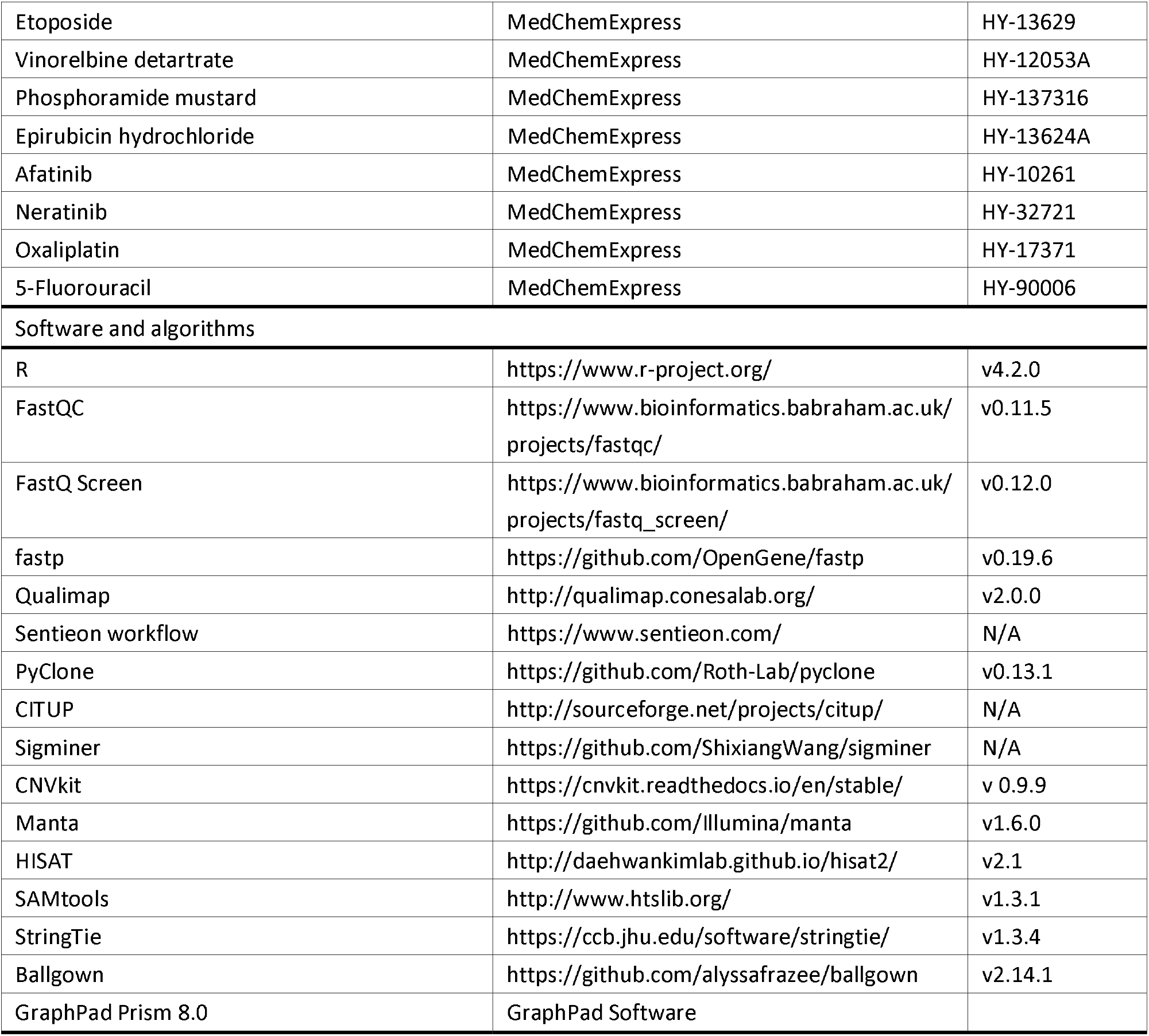

## Resource availability

### Lead contact

Further information and requests for resources and reagents should be directed to and will be fulfilled by the Lead Contact, Yuanting Zheng.

## Materials availability

Organoids established in this study are available from the lead contact with a completed Materials Transfer Agreement.

## Data and code availability

The raw sequence data reported in this paper have been deposited in the Genome Sequence Archive (Genomics, Proteomics & Bioinformatics 2021) in National Genomics Data Center (Nucleic Acids Res 2022), China National Center for Bioinformation / Beijing Institute of Genomics, Chinese Academy of Sciences (GSA-Human: HRA003940) that are publicly accessible at https://ngdc.cncb.ac.cn/gsa-human.45,46 Any additional information, including the code required to reanalyze the data reported in this work paper is available from the lead contact upon request.

## Experimental model and study participant details

### Patient information

The patient provided written informed consent to participate in the clinical trial, abiding by the principles of the Helsinki Declaration. All biopsies, organoid establishments, and molecular assays were performed with approval from the Medical Ethical Council of Shanghai Chest Hospital (approval number: KS(Y)2078). Briefly, the patient was a 48-year-old woman with multiple metastatic NECC who received three surgeries and adjuvant chemotherapy. The primary lesion was diagnosed as NECC combined with adenosquamous carcinoma. The two lung metastasis tissues diagnosed as NECC were established into two organoid lines, consecutively.

## Method details

### Organoid establishment and culture

The two lung metastatic tumor tissues resected by surgery were stored in ice-cold 1× Advanced DMEM/F12 (Gibco) containing antibiotics in conical tubes and processed for organoid culture within 24 hours. Briefly, non-epithelial components in tumor tissues were removed using surgical scissors or scalpels. Then the epithelial components were rinsed with DPBS twice. The tumor tissues were minced into cubic piece of 1-3 mm^3^ in a 6-cm cell culture dish using surgical scissors or scalpels and digested for 1 h using 400 U/mL collagenase I (Thermo), 10 U/mL dispase type I (Roche), 10 μM Y-27632 (OrganRegen) and 1:500 Primocin (Invivogen) in Advanced DMEM/F12 (Gibco) in a 15-mL conical tube at 37°C. Mixture was pipetted vigorously every 15 min until the contents could go through the 5-mL pipette tips. The digestion reaction was terminated by adding FBS to a final concentration of 2% and then filtered through a 100-μm cell strainer. The filtered cells were centrifuged and collected at 200 g for 3 min at 4℃. In case of a visible red pellet, the pellet was resuspended in 2 mL of Red Blood Cell Lysis Solution (Solarbio) at 37℃ for 3 min to lyse the erythrocytes. Then the pellet was washed with Advanced DMEM/F12 twice and resuspended in Matrigel at a density of approximately 10,000 cells per 25 μL. The mixture was plated on the bottom of a 24-well cell culture plate in drops of ∼25 μL each. The culture plate was placed into a humidified incubator at 37℃ and 5% (vol/vol) CO_2_ for 20 min to let the Matrigel solidify. Once the Matrigel drops were solidified, cells were overlaid with cervical cancer organoid culture medium, composed of 1× Advanced DMEM/F12, 100 U/mL Pencillin/Streptomycin (Gibco), 1× GlutaMax (Gibco), 10mM HEPES (Gibco), 1× B27 supplement (Gibco), 1.25 mM N-Acetylcysteine (Sigma), 5 mM nicotinamide (Sigma), 500 ng/mL human recombinant R-spondin1 (OrganRegen), 100 ng/mL human recombinant Noggin (OrganRegen), 25 ng/mL human recombinant FGF-7 (OrganRegen), 100 ng/mL human recombinant FGF-10 (OrganRegen), 50 ng/mL EGF (OrganRegen), 500 nM A83-01 (OrganRegen), 500 nM SB202190 (OrganRegen) and 5 μM Y-27632 (OrganRegen). Organoids formed in approximately two weeks. After that, organoids were passaged every 1-2 weeks by incubating organoids for 5-10 min in TrypLE Express (Gibco) at 37℃ and mechanical disruption to make organoids fall apart, followed by washing and replating in Matrigel. Organoids were cryopreserved in Organoid Cryopreservation Medium (bioGenous) as master and working biobanks. Organoids less than passage 10 were used in experiments.

### Immunofluorescent staining

The cervical tumor tissues were fixed in 4% paraformaldehyde (PFA) at 4℃ for 48 h and then dehydrated and embedded in paraffin. To prepare organoids for pathological identification, organoids cultured in Matrigel were washed and collected to the bottom of tubes, then were fixed in 4% PFA for 2 h. After that, organoids were washed by 1× PBS and resuspended by 50 μL 3% warm agarose, followed by dehydration and embedding in paraffin. Sections were cut and hydrated before staining. The standard protocols for H&E staining and immunofluorescent staining were used. For immunostaining, samples were incubated with primary antibodies (see Key Resources Table) overnight and subsequent incubated with secondary antibodies labeled with Alexa Fluor 488- (Jackson ImmunoResearch, 115-545-003, 3 μg/mL) or Cy3- (Jackson ImmunoResearch, 111-165-003, 6 μg/mL) for 1h. Cell nuclei were counter-stained with DAPI (ASGB-BIO). Images were taken by cellSens system of Olympus microscope or a confocal microscope (OLYMPUS, FV3000).

### Organoid growth curve

Organoids were collected and gently grounded through a 70-μm strainer twice with a 2.5-mL syringe plunger. The suspension was mechanical disrupted and filtered through a 40-μm strainer to ensure the organoids fall apart into single cells. Cells were washed in Advanced DMEM/F12 and counted using haemocytometer. 10,000 cells were plated per 3 μL Matrigel drop into 96-well plate and overlayed with 100 μL of medium. For the growth curve, the cell viability was measured with cell counting kit-8 (CCK8) assay in week 0, week 1, week 2, and week 3 after seeding. The experiments were performed in three biological replicates.

### In vitro drug sensitivity assay

Organoids were collected and gently grounded through a 70-μm strainer twice with a 2.5 mL syringe plunger. Next, the cell clusters in the filtrate were measured and resuspended in Matrigel in a density of 10000 cells/μL, then 3 μL drops of the Matrigel cell cluster suspension were dispensed into the 96-well plates, followed by adding 10 mL culture medium each plate. Seven days after plating the cells, different drugs were added at concentrations of 6 doses. Seven days after adding the drugs, the cell viability was measured by using CCK8 Assay (APExBIO). Results were normalized to vehicles (100%) and baseline control (w/o adding culture medium at day 7). Images of each well were screened with the Operetta CLS High Content Analysis System (PerkinElmer) on day 7 and day 14. The experiments were performed in 3 biological replicates.

### Blood concentration data collection for phase I clinical trials of drugs

The range of C_max_ for each drug was obtained by searching PubMed database, using the keywords “pharmacokinetics” and “phase I clinical trials”. Trials on combination drugs were excluded and papers on phase I clinical trials of single drugs were selected. C_max_ for each drug was obtained from the selected representative published paper and combined with the data from DrugBank Online (https://go.drugbank.com/). When the IC50 is below the lower limit of C_max_, the drug is defined as a sensitive drug; when the IC50 is above the upper limit of C_max_, the drug is defined as a tolerated drug.

### DNA and RNA extractions

Genomic DNA of patient-derived organoids and matched WBC were isolated using SDS (Thermo Fisher, USA) method following the recommended instructions. Genomic DNA from FFPE tissues were extracted using TIANGEN FFPE DNA Kit (cat#DP330). The degradation and contamination of DNA was monitored on 1% agarose gel. The concentration of DNA was measured using the Qubit® DNA Assay Kit in the Qubit® 3.0 Flurometer (Invitrogen, USA). For total RNA extraction, the organoids were processed with TRIzol (Invitrogen, USA) according to the manufacture instructions. The total amount and integrity of RNA were assessed using the RNA Nano 6000 Assay Kit of the Bioanalyzer 2100 system (Agilent Technologies, CA, USA).

### Whole-genome sequencing

The WGS library was generated using NEB Next® Ultra™ DNA Library Prep Kit (NEB, USA), following the manufacturer’s recommended protocol. Two technical replicates were performed in parallel for each sample. DNA library concentration was measured by Qubit®3.0 Flurometer (Invitrogen, USA). Library size distribution was analyzed using NGS3K/Caliper and quantified using real-time PCR (3nM). The libraries were sequenced with NovaSeq 6000 with paired-end 150-bp read-length in Novogene Company (Beijing, China). FastQC v0.11.5 (https://www.bioinformatics.babraham.ac.uk/pro-jects/fastqc/), FastQ Screen v0.12.0 and Qualimap v2.0.0 were using for data quality control.^47,48^ Sequences were mapped to GRCh38 (https://gdc.cancer.gov/about-data/gdc-data-processing/gdc-reference-files) using BWA-mem. vcf files were generated from fastq files using Sentieon Genomics software (https://www.sentieon.com/). The Sentieon workflow includes mapping reads to reference, calculating data metrics, removing duplicates, Indel realignment, base quality score recalibration (BQSR) and performing variant calling with TNseq. Default settings were used for all processes.

### Somatic single nucleotide variants

Somatic SNVs and indels were detected by Sentieon TNseq. High confidence SNVs and indels were filtered by the following steps: 1) all sites were in the overlap of the two technical replicates; 2) depth of the SNV/indel site is more than 10; 3) variant allele fraction of all sites is more than 5% 4) SNVs and indels within the exonic region yield the preliminary dataset. TMB was defined as the number of somatic mutations in the coding region per megabase, including SNVs and indels.

PyClone was used to cluster the final meaningful SNVs/indels.^49^ PyClone is a Bayesian clustering method to estimate the cellular prevalence and account for allelic imbalances. The program of ‘run_analysis_pipeline’ in PyClone was conducted with default settings. The clonal phylogeny was built using CITUP.^50^

Sigminer was used for single base substitution (SBS) mutation signature analysis.^51^ Sigminer is an one-stop solution for de novo signature extraction, reference signature fitting, signature stability analysis, sample clustering. De novo mutational signatures were compared to curated signatures in COSMIC using cosine similarity. SBSs with the highest match scores were selected.

### Somatic copy number alterations (SCNAs)

CNVkit v 0.9.9 was performed with default parameter on paired tumour-normal sequencing data to get genome-wide somatic copy number alterations.^52^ The intersection of the two technical replicates from each sample is taken as the set of high-confidence CNVs. The gain is the part whose log2 ratio of copy number relative to normal sample was more than 0.32; the loss is less than -0.41. Cancer related genes were select and were compared with the RNA-seq data. The cancer genes are from TCGA and COSMIC.^25^

### Somatic structural variations

Manta (v1.6.0) somatic structural variant calling pipeline was performed with default options with tumor and normal BAM files using GRCh38 (https://gdc.cancer.gov/about-data/gdc-data-processing/gdc-reference-files) as the reference genome.^53^ The intersection of the two technical replicates from each sample is taken as the set of high-confidence SVs.

### RNA sequencing

The RNA library was constructed using NEB Next ® Ultra™ RNA Library Prep Kit (NEB, USA), following manufacturer’s recommended protocol. Two technical replicates were performed in parallel for each sample. The library was initially quantified by Qubit2.0 Fluorometer, then the insert size of the library was detected by Agilent 2100 bioanalyzer. After insert size meets the expectation, qRT-PCR was used to accurately quantify the effective concentration of the library (the effective concentration of the library was higher than that of 2nM) to ensure the quality of the library. The mRNA libraries were sequenced with NovaSeq 6000 with paired-end 150-bp read-length in Novogene Company (Beijing, China). Initial processing of the raw fastq reads was carried out using fastp v0.19.6 to remove adapter sequences.^54^ FastQC v0.11.5 (https://www.bioinformatics.babraham.ac.uk/pro-jects/fastqc/), FastQ Screen v0.12.0 and Qualimap v2.0.0 were using for data quality control were using for data quality control.^47,48^ Reads alignment and quantification was conducted using HISAT v2.1, SAMtools v1.3.1, StringTie v1.3.4 and Ballgown v2.14.1.^55-57^ Genome Reference Consortium human genome build 38 (Genome version: GRCh38_snp_tran) and gene model from Ensembl (version: Homo_sapiens.GRCh38.93.gtf) were used for read mapping and gene quantification.

### Gene expression data

The mean value of expression data of the two technical replicates from each sample was taken as the final expression data. Six cervix normal sample expression data were download from the GTEx database; Three cervical paracervical samples was taken as a control, whose expression data were download from TCGA.

### Quantification and statistical analysis

For drug testing shown in figures, statistical analyses were performed with Excel and GraphPad Prism (GraphPad, San Diego, CA, USA). Each drug was diluted with culture medium and dispensed into 96-well plates. Three wells were used for each concentration of each drug. Control wells (vehicles and baseline control wells) were placed in every plate, with at least 3 wells for each control. The half-maximal (50%) inhibitory concentration values (IC50) were calculated with GraphPad Prism 5 software by using non-linear regression. For omics data analysis, all statistical analyses and figure plotting were performed using R version 4.2.0.

## SUPPLEMENTAL INFORMATION TITLES AND LEGENDS

**Figure S1.**
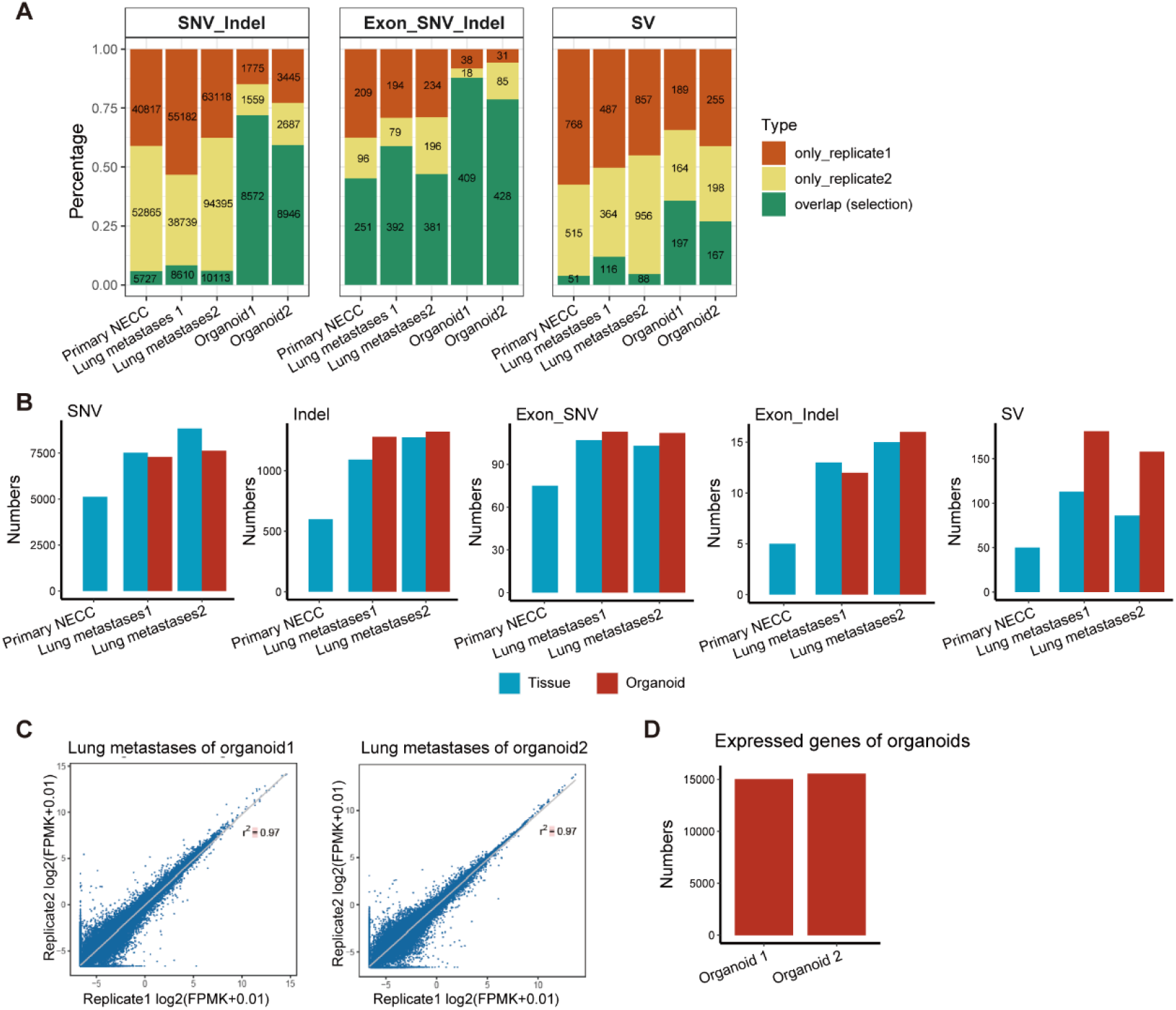
Summary information of WGS and RNA-seq data. (A) Concordance of two technical replicates of WGS. The number marked represents the specific number of each type. (B) The number of SNVs (single nucleotide variation) and Indels on a whole-genome-wide scale; the number of SNVs and Indels on a whole exon genome-wide scale; the number of SVs (structure variations) on a whole-genome-wide scale. (C) Concordance of the two technical replicates of RNA-seq. (D) The total number of expressed genes (FPEK>0.05).

**Figure S2.**
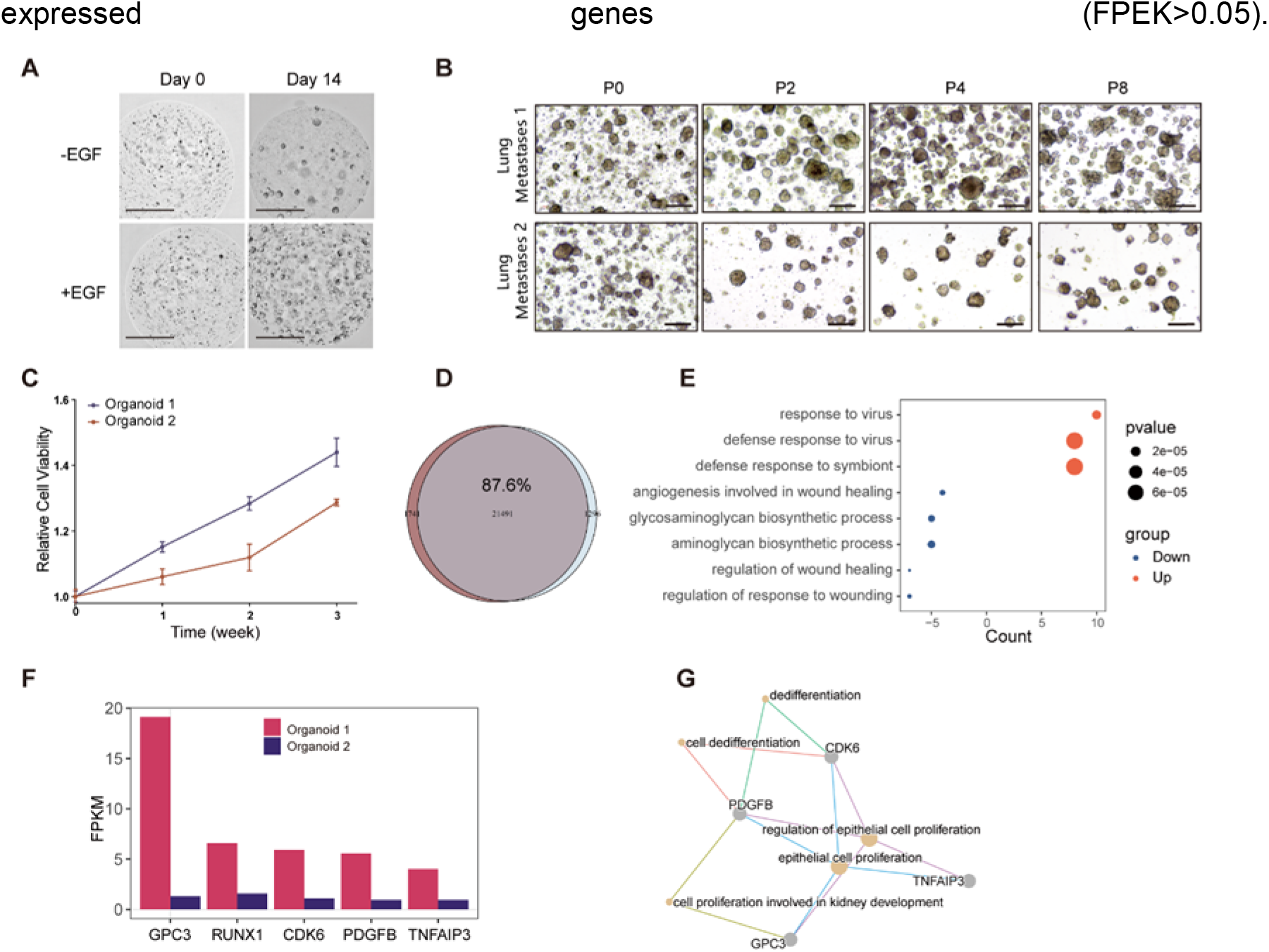
The growth state and RNA expression of the two patient-derived metastatic NECC organoid lines. (A) Growth status of organoids with and without EGF. (B) Growth status of different passages of the two metastatic NECC organoids. (C) Relative cell viability of the two organoids cultured over the course of 3 weeks. (D) Venn plot of the expressed genes of the two organoids. (E) GO analysis; up-regulated genes indicate log (FPKMorganoid2 / FPKMorganoid1) > 2; down-regulated genes indicate log (FPKMorganoid2 / FPKMorganoid1) < -2. (F) Bar plot of FPKM value of five significantly upregulated cancer genes (log (FPKMorganoid2 / FPKMorganoid1) > 2) (G) GO analysis of the five cancer genes; up-regulated cancer genes indicate log (FPKMorganoid2 / FPKMorganoid1) > 2; up-regulated cancer genes indicate log (FPKMorganoid2 / FPKMorganoid1) < -2.

**Figure S3.**
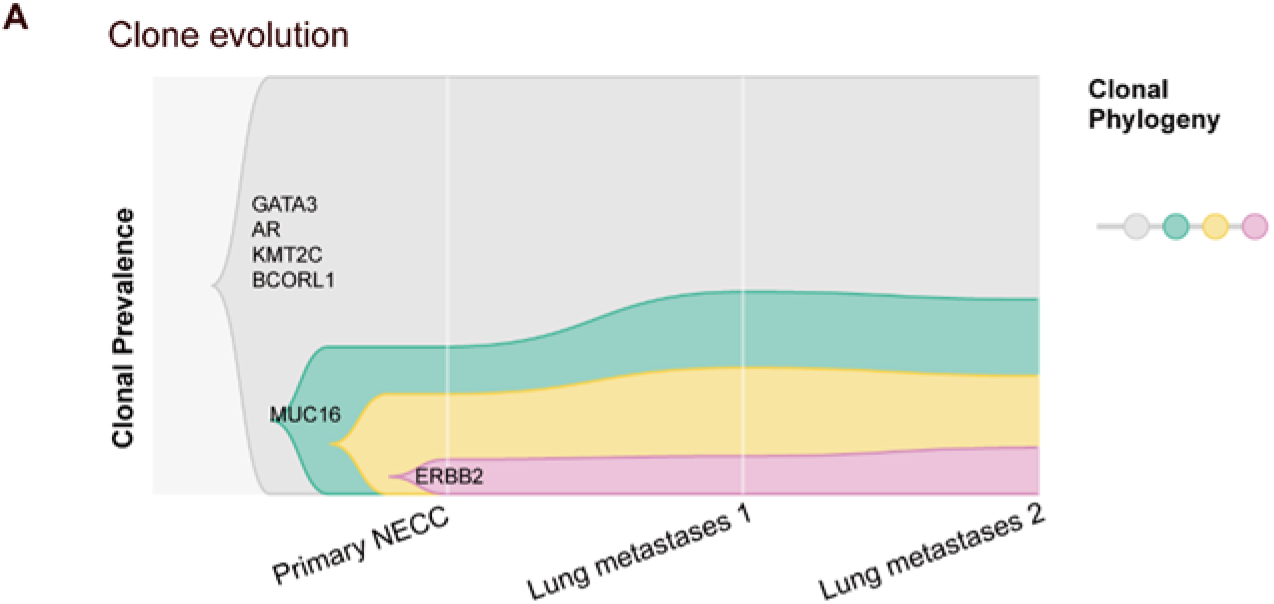
Clone evolution of SNVs/Indels from primary NECC to lung metastases. The y axis denotes the proportion of tumor cells in each clone.

**Figure S4.**
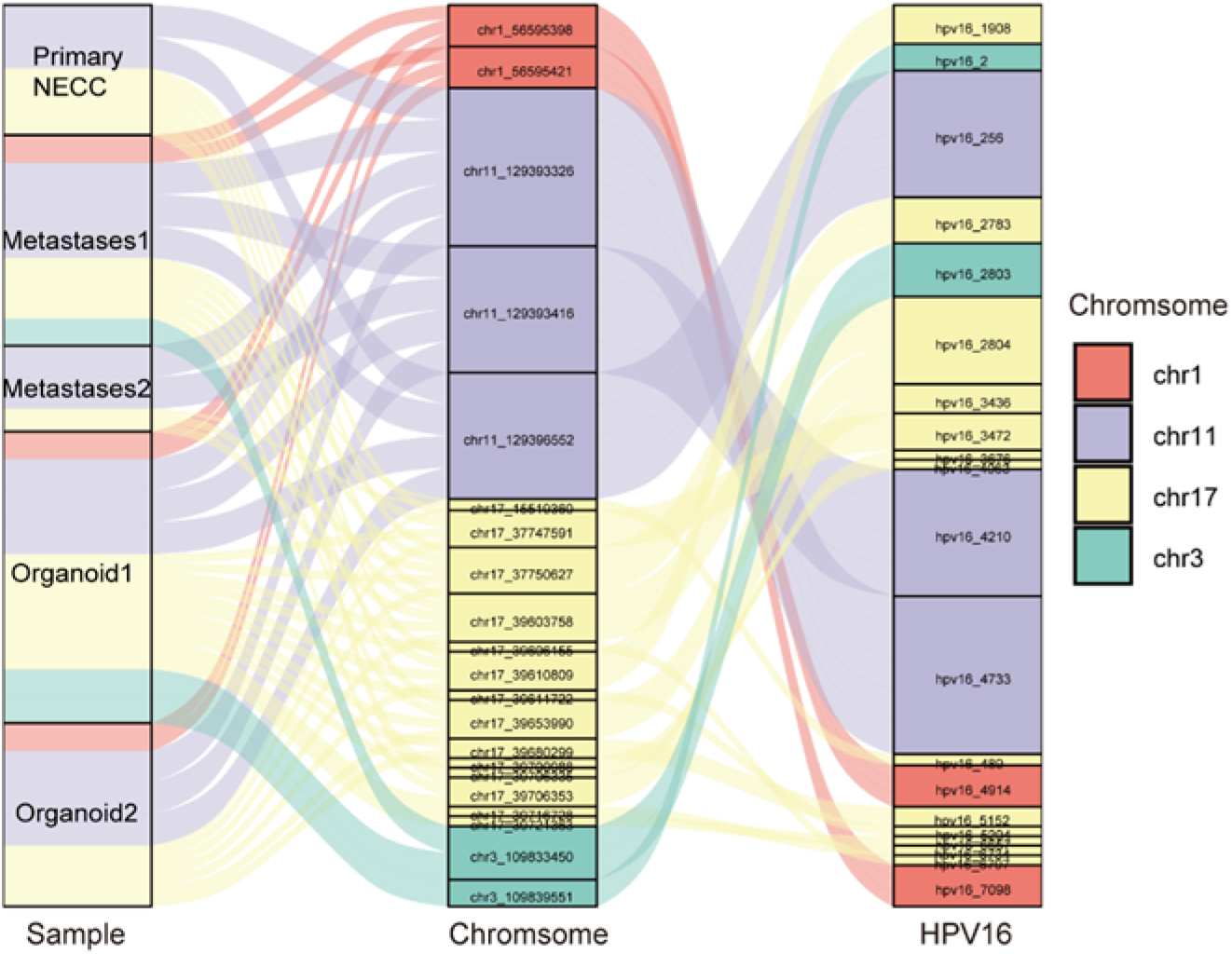
Profiles of HPV16 insertion sites.

**Table S1. The single-nucleotide variant of tumor tissue and corresponding organoid**

**Table S2. The Clonal evolution clusters**

**Table S3. The copy number variation of tumor tissue and corresponding organoid Table S4. HPV16 insertion sites**

**Table S5. mRNA expression [log2(TPM+1)]**

**Table S6. Culture system**

